# Exploring the Interactive Effects of Seed and Site Limitation on Recruitment Dynamics in *Panicum hallii*

**DOI:** 10.1101/2023.06.12.544673

**Authors:** Samsad Razzaque, Thomas E Juenger

## Abstract

Seedling recruitment is a critical life history stage that is likely driven by the interplay of seed and microsite limitation and their interaction with seed traits. Here, we performed a factorial experiment involving seed addition and surface soil disturbance to explore the combined effects of seed and site availability using genotypes characterized by varying seed mass and dormancy traits. Additionally, we included hybrids that exhibited recombined seed trait relationships compared to natural genotypes, allowing us to assess the impact of specific seed traits on establishment across different sites. We detected a significant three-way interaction between seed addition, site conditions, and soil surface disturbance, influencing both seedling and adult recruitment in *Panicum hallii*, a perennial grass found in coastal mesic (lowland) and inland xeric (upland) habitats. This recruitment pattern suggests that lowland and upland establishment at foreign site is constrained by the interplay of seed and site limitations. Notably, soil surface disturbance facilitated recruitment of the upland genotype while constraining recruitment of the lowland genotype across all sites. Our results highlight the importance of seed size and dormancy as key factors impacting recruitment, suggesting a potential interactive relationship between these traits.

## Introduction

Understanding the factors that govern population dynamics in natural populations is a fundamental challenge in the field of population ecology. When a plant species fails to establish successfully at a particular site, the underlying cause can be attributed to either seed or site limitation (Clark et al. 2007). Seed limitation occurs when the density of seeds fails to reach maximum levels at potential recruitment sites within a population Eriksson and Ehrlén 1992a; Svenning and Wright 2005). On the other hand, site limitation arises when the abiotic environment (such as soil characteristics and moisture availability) and/or the biotic environment (including the presence of competitors or seed predators) constraints the processes of germination, growth, and reproduction Nathan and Muller-Landau 2000; Harper 1977; Gross 1980; Bergelson 1990; Hilton and Boyd 1996; Houle 1996; Clark et al. 2007; Beckman and Rogers 2013). Both seed and site limitations have been observed in natural systems Clark et al. 2007; Chave 2004). However, the interplay of various factors, including the evolutionary diversification of seed traits within populations and their interaction with local conditions (both biotic and abiotic), can play a causal role in recruitment dynamics. For instance, the emergence of seedlings may be influenced by the interaction between seed traits (such as size or dormancy) and the heterogeneity of microsite conditions (such as soil moisture), where the germination of dormant seeds depends on the availability of moisture in the soil Limón and Peco 2016; Merino-Martín et al. 2017; Lewandrowski et al. 2017). Nevertheless, viable seeds alone do not guarantee successful recruitment, and the presence of viable seeds is a necessary but insufficient condition for recruitment to occur. Therefore, a comprehensive understanding of species recruitment limitation necessitates a careful investigation that incorporates multiple factors (including seed traits, seed viability, habitat structure, and different life stages) that potentially influence recruitment patterns in diverse habitats. By considering these intricate dynamics, we can gain deeper insights into the complexities of recruitment processes within species.

Theoretical models offer various predictions concerning seed and site limitation from a community ecology perspective. One perspective conceptualizes seedling recruitment as a lottery system, where emergence is solely determined by the density of arriving seeds across species, including both poor competitors and dominant species Chesson and Warner 1981; Warner and Chesson 1985; Cornell and Lawton 1992; Hurtt and Pacala 1995). Conversely, models based on a competition/colonization trade-off suggest that recruitment relies entirely on the competitive ability (superiority/dominance) of a species, with superior competitors establishing wherever seed is present, regardless of the abundance of inferior competitors’ seeds Armstrong 1976; Hastings 1980; Shmida and Ellner 1984; Tilman 1994). Resource limitation, on the other hand, is expected to result from the interplay of factors such as competitive ability, regeneration niches, relative abundance, and the quality of suitable microsites for seed germination (e.g., light, moisture, temperature regimes) Turnbull et al. 2000; Clark et al. 2007; Pearson et al. 2002; Muller-Landau et al. 2002). In cases of site limitation, disturbance may play a crucial role in determining the availability of microsites for seedling establishment by altering competitive interactions, abiotic conditions, selective damage or mortality, or trophic interactions Petraitis et al. 1989; Luisa 2012; Hulme 1994; Belsky 1992; Bazely and Jefferies 1986; Jutila 1997; Coffin et al. 1998; Wootton 1998; Jutila and Grace 2002). While a few empirical studies have explored the interactions between seed and resource limitation Eriksson and Ehrlén 1992a; Juenger and Bergelson 2000b; Seiwa 2000; Uriarte et al. 2010; Aradottir and Halldorsson 2018; Commander et al. 2020), it remains unclear how these patterns are influenced by habitat variation or locally adapted seed traits.

Plants adapted to heterogeneous habitats exhibit distinct strategies in their life history traits. For instance, species adapted to habitats with varying soil water availability often display variations in physiology, development, morphology, and reproductive allocation strategies (Chabot and Bunce 1979; Solbrig et al. 1977; Stebbins Jr 1952; Whittaker 1970; Woodward et al. 2004; Juenger 2013; Roux et al. 2006). Comparative studies suggest that plants adapted to xeric habitats tend to have larger seeds, which provides an advantage in the early stages of seedling establishment by facilitating faster root and shoot growth to acquire ephemeral resources Westoby et al. 1996; Baker 1972; Schimpf 1977; Sorensen and Miles 1978; Stromberg and Patten 1990). Research has also demonstrated that seedling recruitment and establishment are significantly influenced by seed traits and environmental conditions across different habitats Zhang et al. 2010; Phartyal et al. 2020; Zhang et al. 2019). Therefore, variation in seed size plays a crucial role in seedling and adult recruitment in diverse habitats. Moreover, seed size has been found to affect the timing and probability of seedling emergence, seedling survival, reproductive output, and the ability to form persistent seed banks within specific environments Black 1957; Morse and Schmitt 1985; Hammond and Brown 1995; Moles et al. 2005). However, most studies have focused primarily on seed size and observations from a limited number of systems. Further investigation is needed to understand how diverse seed traits interact with the environment, impact plant performance, and influence recruitment Abbas et al. 2021; Saatkamp et al. 2019).

Seedling emergence and survival represent the most vulnerable stage in a plant’s life cycle, subject to intense selection pressure and high mortality rates Donohue et al. 2010; Donohue 2014; Poorter 2007; Kitajima and Fenner 2000; Mercer et al. 2011). Allocation strategies of plant species may vary depending on the habitat or plant community context, leading to phenotypic and genetic trade-offs within species across different habitats. A recent study of the perennial grass *Panicum hallii* revealed a genetic trade-off between seed mass and seed dormancy, with an antagonistic relationship associated with establishment in xeric and mesic adapted genotypes Razzaque and Juenger 2022). Habitat quality and local competitive environments likely play significant roles in shaping the selection pressures and evolution of seed and seedling phenotypes between *P. hallii* ecotypes. We hypothesize that *P. hallii* ecotypes have undergone strong selection pressures resulting in the evolution of either large seeds with high dormancy or small seeds with low dormancy traits along the pronounced abiotic stress/competition gradient within the species’ range. However, few studies have disentangled the effects of putatively functional seed traits (e.g., seed mass or seed dormancy) to explore their individual and combined impacts on recruitment in natural populations. Therefore, it would be valuable to assess the effects of these trait combinations on recruitment. To break specific trait relationships, one approach is to generate recombinant materials with trait combinations opposite to those of the parental genotypes, allowing us to determine the significance of individual traits or specific trait combinations for recruitment.

In this study, we investigate the role of seed and site limitation, as well as the variability in seed size and dormancy traits, in the establishment of *Panicum hallii* across contrasting habitats. *P. hallii* is a highly self-fertilizing species. It is generally found in the southwestern USA and northern Mexico, exhibiting significant morphological divergence between its two ecotypes adapted to xeric (*P. hallii* var. *hallii*) and mesic (*P. hallii* var. *filipes*) habitats, previously classified as distinct varieties or subspecies Gould 1975; Waller 1976; Lowry et al. 2013; Lovell et al. 2018; Palacio-Mejía et al. 2021). The upland ecotype predominantly grows in rocky, calcareous soil, avoiding competition, and exhibits a broader natural distribution range compared to the lowland ecotype based on herbarium records and field observations (Palacio Mejía 2018; Palacio-Mejía et al. 2021). Conversely, the lowland ecotype thrives in mesic coastal areas, facing high levels of competition from numerous perennial and annual species Gould et al. 2018; Lovell et al. 2018; Lowry et al. 2015; Palacio Mejía 2018; Palacio-Mejía et al. 2021). It is worth noting that the two ecotypes coexist in a limited area within their range Waller 1976; Lowry et al. 2013; Palacio-Mejía et al. 2021). Hence, an important aspect to consider regarding *P. hallii* ecotype establishment is the factors constraining their distributions, the infrequent overlap between ecotypes, and the adaptive strategies that have evolved in response to contrasting habitats.

This study aimed to unravel the underlying factors driving the establishment of *P. hallii*, specifically examining the influence of seed limitation and site availability in diverse habitats. By conducting experiments in two distinct field sites representing mesic and xeric environments, we employed plot-based seed addition and soil disturbance techniques to assess their effects on overall seedling and adult recruitment. Additionally, we delved into the role of seed size and dormancy traits by utilizing two natural genotypes and two recombinant lines that possessed opposite combinations of these traits, thus investigating their potential facilitation or constraint on adaptation across varying sites.

## Materials and Methods

### Plant materials, seed collection and experimental design

In our study, we utilized two distinct inbred genotypes representing the upland (*hallii*) and lowland (*filipes*) ecotypes, namely HAL2 and FIL2, respectively. These genotypes were collected from two locations: the upland genotype was obtained from the Lady Bird Johnson Wildflower Center in Austin, TX, USA (30.19°N, 97.87°W), while the lowland genotype was sourced from a natural area associated with the Corpus Christi Botanical Garden in South Texas (27.65°N, 97.40°W). HAL2 and FIL2 have been extensively studied and recognized as representative genotypes for genome assembly and ecotype characterization Lovell et al. 2018).

We generated two recombinant inbred lines (RILs) through hybridization between HAL2 and FIL2. These RILs, designated as RIL1 and RIL2, were derived from the F2 progeny of a single F1 hybrid and underwent seven generations of single seed descent breeding to achieve homozygosity Khasanova et al. 2019).

Typically, the upland ecotype displays larger seeds with higher levels of dormancy, while the lowland ecotype exhibits smaller seeds with lower dormancy. Both seed size and dormancy traits demonstrate significant broad-sense heritability (H^2^ = 0.89 and 0.24 for size and dormancy, respectively) and exhibit minimal plasticity, indicating that these traits are relatively stable and influenced by various genomic regions (QTLs) Razzaque and Juenger 2022) . As each trait is driven by unique genetic factors, the hybrid progeny (RILs) exhibit diverse patterns of trait variation and covariation. Specifically, RIL1 showcases the combination of small seed size and high dormancy, while RIL2 exhibits large seed size and low dormancy (Figure 1). These RILs were selected for further study based on their distinct trait profiles, as previously characterized by Razzaque and Juenger (2022).

**Figure 1:**
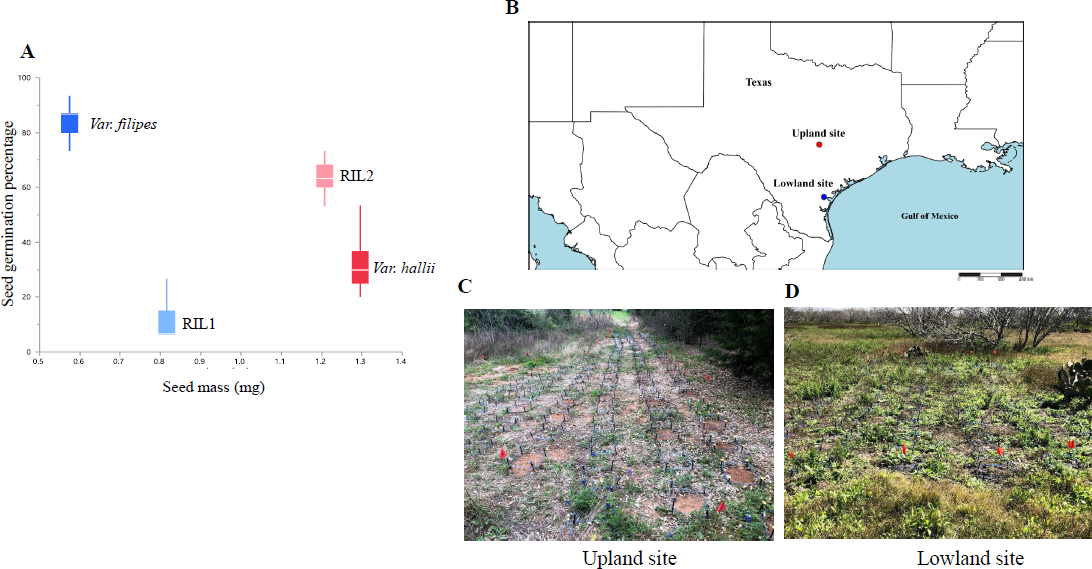
Plant materials and location of experimental sites. A: Plant material used in this experiment included two natural inbred genotypes (FIL2 and HAL2) and two recombinant inbred lines (RIL1 and RIL2). Y-axis represents the seed germination percentage conducted in laboratory condition and X-axis represents the mean seed mass for each line. Each error bar is constructed using ±1 standard error from the mean. B: Geographic location of two experimental sites in Texas representing a typical xeric and mesic habitat of *P. hallii*. C & D: Experimental sites prepared for the seed addition with wires and cones to delimit plots. These photos were taken before seed addition in January 2019.

In April 2018, we obtained seeds for the four selected genotypes by cultivating 30 plants of each genotype in a controlled growth chamber at UT Austin. Initially, the plants were germinated in petri dishes and subsequently transplanted into one-gallon pots filled with a growth medium composed of Promix and Turface at a ratio of 6:2. To ensure optimal growth conditions, the plants were subjected to a 12-hour light and 12-hour dark photoperiod, with daytime temperatures set at 28_J and nighttime temperatures at 24_J. Watering was conducted every three days to maintain adequate soil moisture levels. Seed collection was carried out when the seeds reached maturity, with a target of approximately 100 grams of seeds per genotype. Following collection, the seeds were stored at room temperature for a period of six months to undergo a process known as after-ripening. Our previous observations indicated that freshly collected seeds exhibited considerable variability in germination traits under laboratory conditions. However, by allowing them to undergo after-ripening for around five months, we were able to minimize this variability. Before commencing the seed addition experiment, we conducted germination tests to determine the germination percentage of the collected seeds (Figure 1).

For the reciprocal seed addition experiment, we chose a representative xeric site located at the UT Brackenridge Field Laboratory (30.28°N, 97.78°W); BFL; Austin, Texas) and a representative mesic site located at the Nueces Delta Preserve (27.92°N, 97.61°W); NDP; Odem, Texas, Figure1) on the Gulf coast. At BFL, we find var. *hallii* in dry and low cover grassland communities composed primarily of short-lived annual grasses (e.g., *Avena fatua*, *Bromus catharticus*). In contrast, we find var. *filipes* at the NDP in competitive coastal prairie environments with a high density of perennial grasses (e.g., *Bouteloua rigidiseta, Nassella leucotricha*). We selected these sites and locations for experimental plots near areas with naturally occurring *P. hallii.* We established 0.5m x 0.5m plots for the experiments in early February 2019.

It is important to note that we could not conduct the reciprocal transplant work at the exact sites of origin for our representative lines. Unfortunately, the collection location for the lowland ecotype has been converted to a residential area and the other site does not allow manipulative experiments. As such, we picked nearby sites with populations of *P. hallii* occurring in natural vegetation. The local populations at these sites are genetically and phenotypically similar to the representative lines we studied (Palacio-Meija et al. 2021). Our sites differ in soil, cover vegetation, and climatic features that clearly represent the natural ecoregions (Edwards Plateau Savanna and Western Gulf Coastal Grassland) and ecological preferences of the ecotypes. We also note that our experiment is pseudo-replicated from the perspective of contrasting xeric and mesic habitats.

The experimental design encompassed several factors, including genotype (upland/lowland/two RILs), site type (xeric/mesic), and soil surface disturbance (disturbed/undisturbed) in addition to control plots. To achieve a comprehensive analysis, we employed a fully factorial experiment, with each combination of factors replicated in ten plots per site. In total, we investigated 200 plots measuring 0.5m x 0.5m (consisting of 5 genotypes: 4 genotypes plus 1 unseeded control plot) across two field sites and two disturbance treatments. Prior to seed addition, the plots were randomized into treatment categories.

For the disturbed plots, we meticulously removed all vegetation by cutting the aboveground plant material to ground level and then disturbed the soil using a rake. Control plots were left untouched, representing their natural state. To ensure accurate identification and minimize confusion, we also planted ten HAL2 and ten FIL2 genotypes in one-gallon pots at both field sites. These potted plants served as reference points for identifying seedlings in the field. Starting from the day of seed addition, we conducted regular inspections of both sites at 15-day intervals. During each visit, the plots were thoroughly examined for the emergence of new seedlings. Each newly identified seedling was carefully tagged with a mini rubber band. Once identified and marked, the seedlings were continuously monitored from the early establishment phase until they reached the flowering stage. Flowering served as a metric to determine the count of adult reproductive plants. The raw count data for both seedling and adult plant observations can be found in Supplementary File 1.

### Experimental design for seed longevity test

To assess seed longevity at both the mesic and xeric sites, we conducted a burial experiment. We buried a total of 280 small mesh bags, with 70 bags each containing HAL2 and FIL2 seeds (70 bags x 2 genotypes x 2 sites). The bags were buried at a depth of 30 cm. The burial took place in late summer, specifically on August 25 at the xeric (BFL) site and August 27 at the mesic (NDP) site in 2018. At regular intervals of 30 days after the burial date, we collected the seed bags. Subsequently, we performed germination assays on the collected seeds in a growth chamber. To facilitate germination, we removed the seed coats using a well-established method for forcing germination in *P. hallii*, as described in Razzaque and Juenger (2022) .

The duration of the study spanned 18 months, during which we collected buried seeds from each field location. The seeds obtained from the bags were germinated in petri dishes containing sand and water, following the methods and growth chamber conditions outlined in Razzaque and Juenger (2022). After 20 days from the first day of seedling germination for each trial, we counted the germinated seedlings (see Supplementary File 2). Using this data, we calculated the germination percentage as follows: Germination percentage = (number of seedlings / total number of seeds) * 100.

### Statistical Analyses

To analyze seed bank longevity and explore the factors influencing germination, we employed a factorial model incorporating Genotype (G), Time (T), Site (S), and their interactions. The Genotype factor represented the HAL2 or FIL2 genotypes, the Site factor represented the xeric or mesic sites, and the Time factor accounted for the sampling time points (18 in total). To assess the relationship between these factors and seed germination, we employed a generalized linear model (GLM) with a Poisson distribution and log link. Specifically, we analyzed the total number of germinated seedlings per genotype. Significant effects observed for Genotype (G) would indicate genotype-related differences in seed bank persistence. Significant effects for Time (T) would suggest temporal influences on seed bank persistence, while significant effects for Site (S) would indicate site-specific influences on seed bank persistence. Moreover, significant interactions between genotype and sites or time would suggest genetic variation in plasticity regarding seed bank persistence. To evaluate the model’s goodness of fit, we utilized generalized r-square statistics, which provided an indication of how well the model captured the observed data.

To gain insights into the factors influencing population dynamics between the lowland and upland genotypes of *P. hallii*, we conducted analyses on seedling and adult recruitment, considering the effects of seed addition (SA), Site (S), Disturbance (D), and their interactions. To quantify the impact of these factors on recruitment, we determined the final counts of unique seedlings and reproducing adult plants. Subsequently, we employed a generalized linear model (GLM) with a Poisson distribution and log link function to examine the relationship. Seed Addition (SA) encompassed both control plots without seed addition and plots with seed addition. Site (S) represented the xeric (BFL) and mesic (NDP) sites, while Disturbance (D) accounted for plots with soil surface disturbances versus undisturbed control plots. In the final model, we assessed the effects of SA, S, and D separately and also explored their potential interactive effects on seedling and adult recruitment. To evaluate the goodness of fit, we employed Pearson goodness-of-fit statistics and concluded that the Poisson GLM adequately captured the observed data.

The complexity of our full model, including higher order interactions, presents challenges in interpreting the factors influencing seedling and adult recruitment in different sites. Therefore, to aid interpretation of the interactions we discovered, we employed a series of contrasts that focused on specific hypotheses derived from the interactions of our manipulated factors and their impact on recruitment. Initially, we compared the control plots with the two natural genotypes, HAL2 and FIL2, in contrasting habitats. This comparison aimed to determine whether there are significant differences indicating whether the population dynamics of genotypes are influenced primarily by seed limitation or site limitation in these distinct habitats.

In order to specifically examine the impacts of disturbance independent of other factors, we conducted separate analyses for HAL2 and FIL2 in response to disturbance. Firstly, we compared FIL2 seedling and adult recruitment while considering the presence or absence of disturbance effects, averaging over the other experimental factors. Subsequently, we performed a similar comparison for HAL2 recruitment. By analyzing the data in this manner, we aimed to identify any significant differences that would suggest whether the manipulation of disturbance favored the recruitment of distinct genotypes across the sites. This approach allowed us to isolate and assess the specific effects of disturbance on the recruitment dynamics of HAL2 and FIL2.

In *P. hallii*, the intertwined nature of seed size and seed dormancy poses challenges in understanding their individual impacts on recruitment. To untangle the effects of these distinct traits, we conducted a series of contrasts analyses. Initially, we compared the two genotypes with small seeds (FIL2, RIL1) against the two genotypes with large seeds (HAL2, RIL2) in diverse habitats. A significant difference observed between these groups would suggest that seed size plays a crucial role in determining recruitment, irrespective of seed dormancy. Subsequently, we compared the two genotypes with high dormancy (HAL2, RIL1) against the two genotypes with low dormancy (FIL2, RIL2) in contrasting habitats. A significant difference between these groups would indicate that seed dormancy is a key determinant of recruitment, regardless of seed size. By conducting these specific contrasts, we aimed to disentangle the influence of seed size and seed dormancy on the recruitment dynamics of *P. hallii*, shedding light on the individual contributions of these traits in differentiating their impact on successful establishment.

Furthermore, we investigated the influence of soil surface disturbance on recruitment by conducting contrast analyses to examine the three-way interactions involving seed size and dormancy. To accomplish this, we initially compared the small-seeded genotypes (FIL2, RIL1) with the large-seeded genotypes (HAL2, RIL2) specifically in disturbed plots. Subsequently, we compared the high dormancy genotypes (HAL2, RIL1) with the low dormancy genotypes (FIL2, RIL2) also in disturbed plots. These planned contrasts allowed us to generate additional hypotheses beyond our overall models. To ensure robust statistical analysis, we applied a rigorous Bonferroni correction for multiple test correction, maintaining a stringent level of significance.

The statistical analysis for this study utilized the JMP Pro 15 software packages from SAS Institute. The visualization of data was accomplished through the utilization of the "graph builder" functions.

## Results

### Seed bank longevity varied significantly between genotypes

We observed a significant interaction between genotype and time (G*T, P < 0.0001) on percent germination, suggesting distinct genetic variations in the persistence of seeds in the soil (Supplementary Table 1). Notably, the upland genotype exhibited a 61% longer seed longevity compared to the lowland genotype across both sites (HAL2 germination (%) = 70.1 ± 2.76, FIL2 germination (%) = 26.8 ± 3.53). The germination rate of the lowland genotype decreased significantly after three months of seed burial, while the upland genotype maintained higher persistence and germination for up to eleven months of seed burial (Figure 2). These findings highlight the limited plasticity but substantial genetic divergence in seed bank persistence between the two ecotypes. This suggests that the upland ecotype may harbor a more substantial and longer-lived seed bank compared to the coastal lowland ecotype.

**Figure 2:**
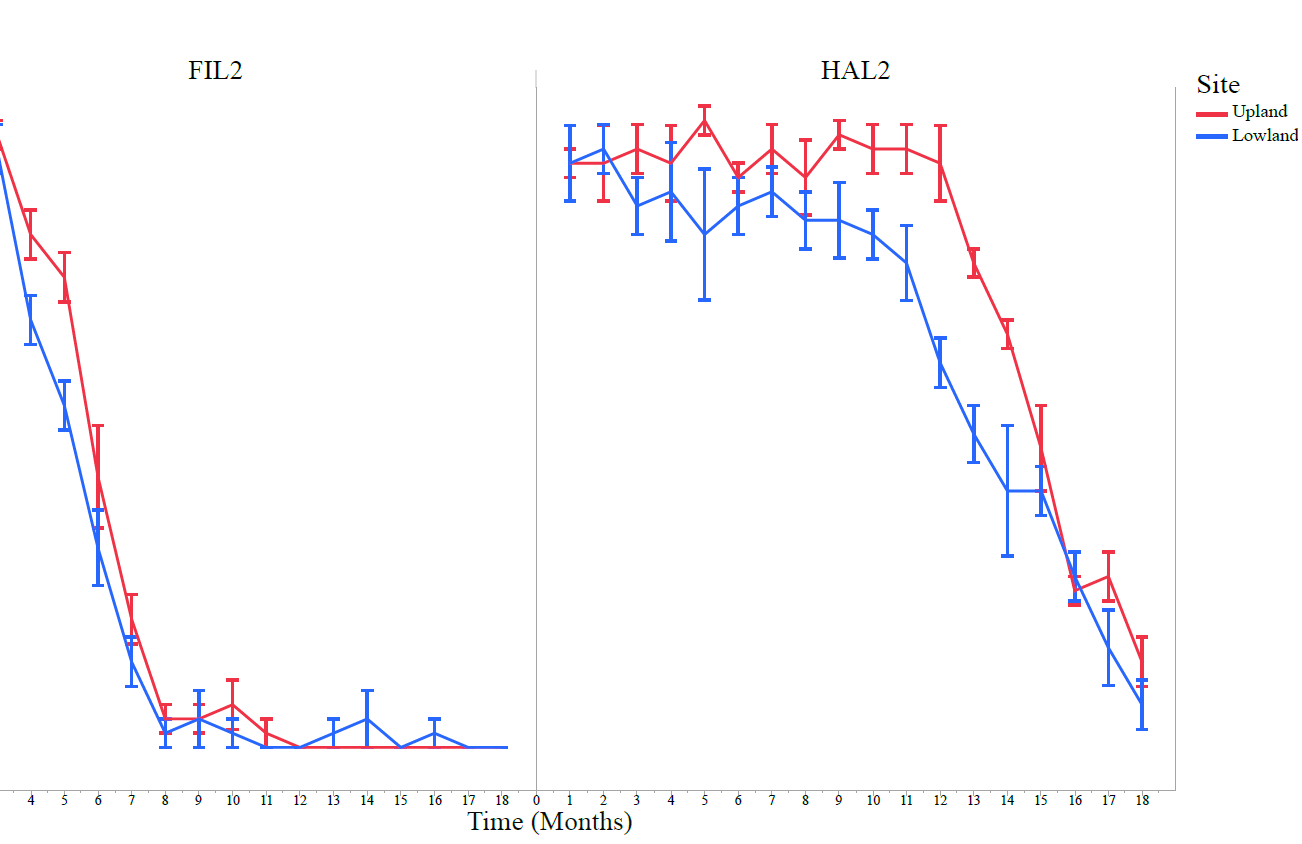
Seed bank longevity between lowland and upland genotypes in contrasting habitats. Lowland (FIL2) and upland (HAL2) genotypes have been tested for seed bank longevity by seed burial experiments in contrasting habitats. The total germinated seedlings of the seeds after burial are plotted on the Y-axis and the time of germination assay in months is shown in X-axis. Both genotypes are plotted separately, the left panel shows the response of the FIL2 genotype, and the right panel shows the response of the HAL2 genotype. The response of mesic (NDP) and xeric (BFL) habitats are colored blue and red respectively. Each error bar is constructed using ±1 standard error from the mean.

### Lowland and upland genotypes are both seed and site limited in natural conditions

We discovered a significant three-way (SA*S*D) interaction for seedling (P < 0.0001) and adult (P = 0.0003) recruitment, as shown in Table 1. The results indicated that seed addition had a notable impact on both seedling and adult plant recruitment across the two sites. Interestingly, our control plots exhibited minimal seedling and adult recruitment, suggesting a limited presence of natural seed banks in the experimental areas. Furthermore, the pattern of the three-way interaction highlighted the significance of multiple factors, including genotype (potentially influenced by seed size and seed dormancy variation) and soil surface disturbance, in shaping the outcomes of seedling and adult recruitment.

**Table 1:**
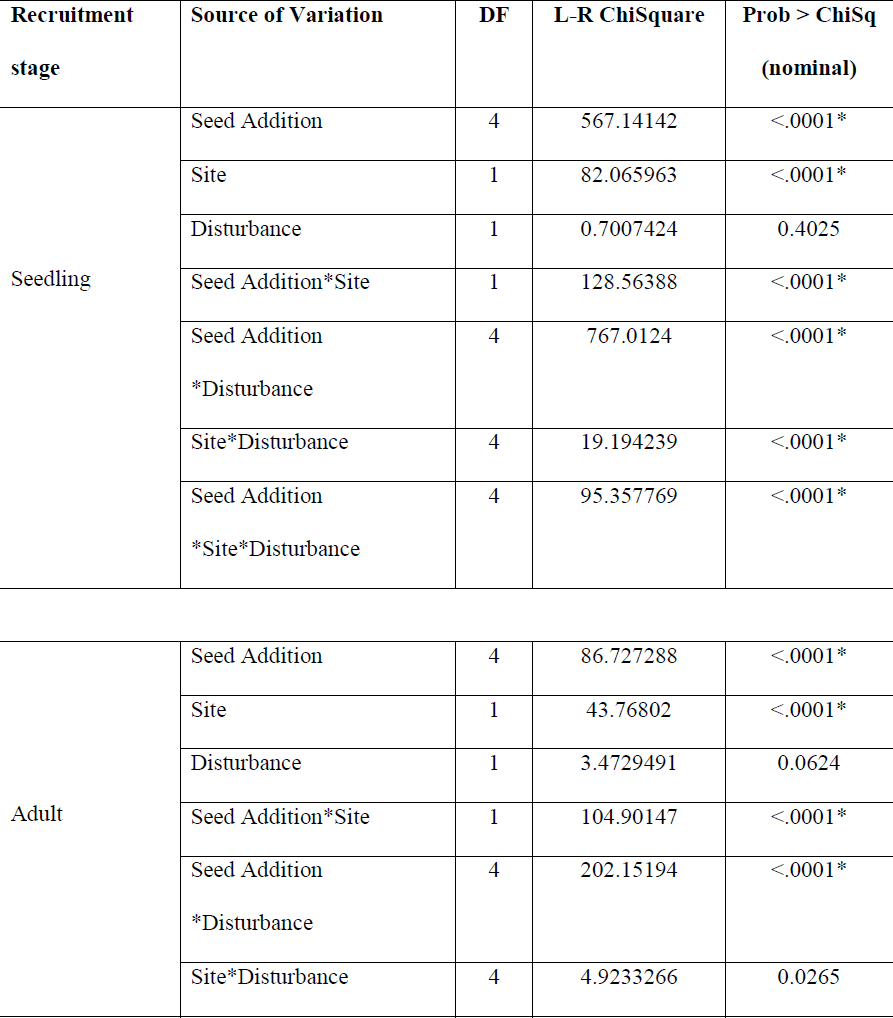
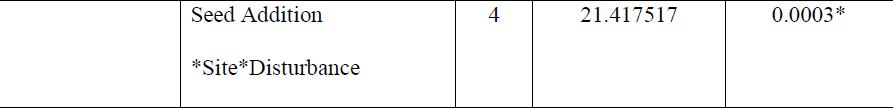
Results of generalized linear model on seedling and adult recruitment in contrasting habitats including disturbed and undisturbed plots. In the factorial model, Genotype, Site and Treatment (seed addition) effect and their interactions have been tested and described in the table. DF = Degrees of Freedom: L-R ChiSquare = Likelihood Ratio Chi Square. ‘*’ Indicates the significance after Bonferroni adjustment at P <0.05.

Due to the complexity of the significant full model with higher order interactions, we used planned contrasts to gain insights into the specific factors influencing *P. hallii* recruitments and to untangle the effects of interactions. Firstly, we compared the seed addition plots with control plots to examine if genotypes derived from lowland and upland ecotypes were seed limited. We observed a significant difference in seedling recruitment (P < 0.0001) when averaged over other experimental factors, indicating that seed addition significantly increased seedling recruitment in the natural plots. Overall, plots that received seed addition exhibited a 91% higher seedling recruitment compared to the control plots. Furthermore, we observed variation in recruitment among genotypes. Specifically, there were significantly more seedlings (∼2.1 times) from FIL2 (lowland) seeds compared to HAL2 (upland) seeds across locations. Notably, FIL2 exhibited similar levels of seedling recruitment in both habitats (NDP (mesic site) = 27.8 ± 3.54 and BFL (xeric site) = 30.6 ± 5.9), whereas HAL2 showed a four-fold greater recruitment at BFL (22.1 ± 2.51) compared to NDP (5.05 ± 0.993) (Figure 3).

**Figure 3:**
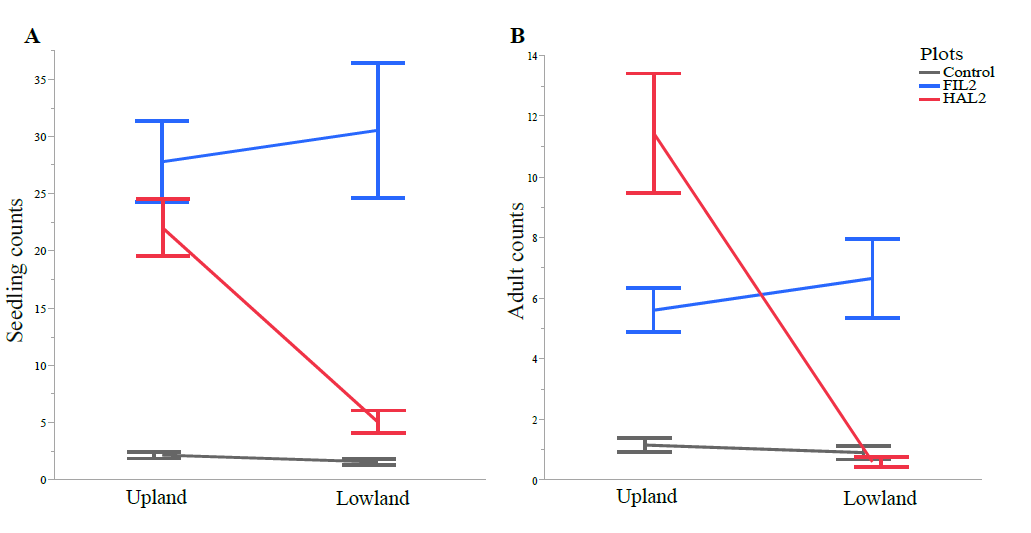
Seedling and adult recruitments in contrasting habitats. For both panels, FIL2 and HAL2 are colored as blue and red, respectively. The control plot is marked grey. The Y-axis shows the number of average recruitments per plot, while the X-axis refers to the site. A. seedling recruitment and B. adult recruitment.

We also observed a significant difference (P < 0.0001) in adult recruitment between seed addition plots and control plots. Seed addition led to an 83% increase in adult recruitment compared to control plots across different habitats. However, overall, the recruitment to the adult stage was relatively low, particularly for HAL2 (upland genotype) seeds sown at the coastal (foreign) site (BFL (xeric site) = 11.5 ± 1.97, NDP (mesic site) = 0.6 ± 0.169). In contrast, FIL2 (lowland genotype) showed consistent adult recruitment across sites (BFL = 5.6 ± 0.731, NDP = 6.65 ± 1.29) (Figure 3). These findings suggest that the genotypes are limited by both seed availability and the specific site conditions under natural circumstances, particularly evident in the limited adult recruitment of HAL2 at the foreign site.

### Disturbance significantly affected establishment in contrasting sites

We observed a significant effect of the disturbance treatment (P < 0.0001) on seedling recruitment for both HAL2 and FIL2 genotypes (Supplementary Table 2). The overall impact of disturbance varied for each genotype across sites. FIL2 seedling recruitment was significantly higher (80% increase) in seed addition plots without disturbance. In contrast, HAL2 recruitment benefited from disturbance, showing a 66% increase in recruits across sites. The pattern was consistent for adult recruitment, with disturbance favoring HAL2 and a lack of disturbance favoring FIL2. HAL2 exhibited an 83% increase in adult recruitment in disturbed plots, while no recruitment was observed in seed addition-undisturbed plots at the NDP site (Figure 4). Notably, HAL2 adult recruitment at the NDP site primarily occurred in disturbed plots. On the other hand, FIL2 had a 79% higher recruitment in undisturbed plots across habitats. These findings suggest that the successful establishment of the HAL2 genotype at the NDP site strongly depends on the effect of disturbance treatments.

**Figure 4:**
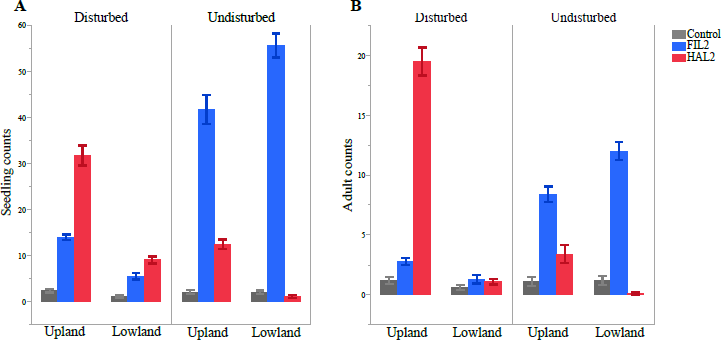
Seedling and adult recruitment in disturbed and undisturbed plots based on the means averaged over the other experimental factors in contrasting habitats. The Y-axis of both panels represents the average number of recruitments and X-axis shows the Sites of the experiments. A. Seedling recruitments under disturbed and undisturbed plots. B. Recruitments of adult plants per plot. Each error bar in the bar chart is constructed using ±1 standard error from the mean.

### Seed sizes significantly interacted with site and disturbance treatment for recruitment

We found a significant difference (P < 0.001) in both seedling and adult recruitment between genotypes with small and large seeds in both habitats (Figure 5, Supplementary Table 2). This indicates that seed size plays a crucial role in determining recruitment, regardless of seed dormancy (Supplementary Table 2). At the BFL site, large-seeded plots exhibited a favorable outcome for both seedling and adult recruitment, showing 13% more seedlings and 57% more adult recruitment compared to small-seeded plots. In contrast, at the NDP site, small-seeded plots were favored, with 70% more seedlings and 68% more adult recruitment compared to large-seeded plots (Figure 5, Supplementary Table 2).

**Figure 5:**
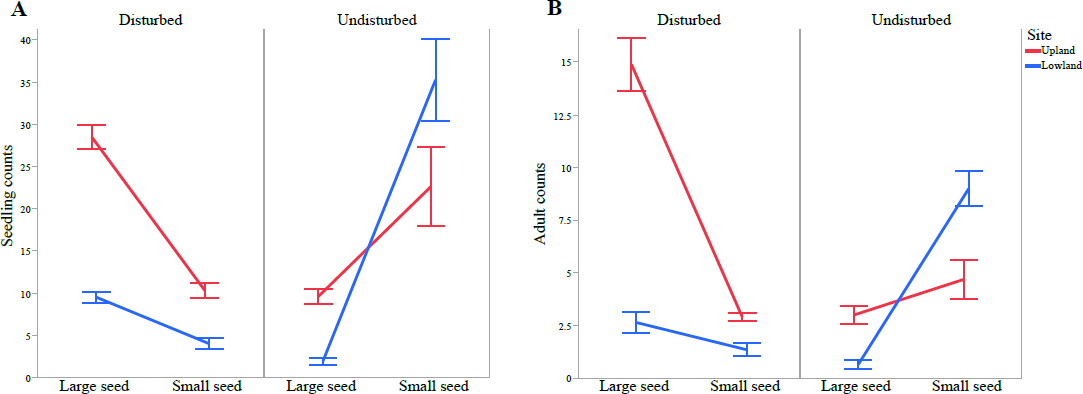
Seedling and adult recruitment for genotypes with big seeds vs genotypes with small seeds group under disturbance treatment conditions in contrasting habitats. The color combination was chosen based on site and treatment effects. The xeric BFL site was colored dark red and the mesic NDP site was colored dark blue. The top panel separated both disturbed and undisturbed plots. The Y-axis represents the mean recruitment counts per plot and X-axis represents seed groups (big seed vs small seed). Each error bar in the bar chart is constructed using ±1 standard error from the mean. A and B: seedling and adult recruitment in contrasting habitats under disturbance treatment conditions.

Furthermore, we observed significant differences in seedling recruitment between large and small-seeded plots in both disturbed and undisturbed conditions across sites. In undisturbed plots, small-seeded plots had 57% and 94% more seedling recruitment compared to large-seeded plots at the BFL and NDP sites, respectively. Conversely, in disturbed plots, large-seeded plots had 63% and 57% more seedlings recruited at the BFL and NDP sites, respectively, compared to small-seeded plots (Figure 5, Supplementary Table 2).

The impact of disturbance on adult recruitment varied across sites. At the BFL site, disturbance favored adult recruitment in large-seeded plots, resulting in 80% more recruitment compared to small-seeded plots. In contrast, at the NDP site, undisturbed plots showed a 92% higher recruitment of adults in small-seeded plots. These differences between seed size groups were statistically significant (P < 0.0001) in both cases. However, there were no significant differences (P = 0.9042) in adult recruitment between large and small-seeded groups in undisturbed plots at the BFL site. Similarly, disturbances did not have a significant effect (P = 0.0693) on adult recruitment at the NDP site (Figure 5, Supplementary Table 2).

### Seed dormancy significantly impacted recruitment across sites

We observed a significant difference (P < 0.0001) in seedling recruitment between the high dormancy and low dormancy groups at both sites, indicating the importance of seed dormancy in recruitment regardless of other experimental factors. Overall, the low dormancy group exhibited 38% and 62% higher seedling recruitment compared to the high dormancy group at the BFL and NDP sites, respectively. In contrast, for adult recruitment, the high dormancy group had 10% higher recruitment than the low dormancy group at BFL, while the low dormancy group showed a 54% higher adult recruitment compared to the high dormancy group at NDP. These pairwise contrasts were statistically significant at P < 0.05 (Figure 6, Supplementary Table 2).

**Figure 6:**
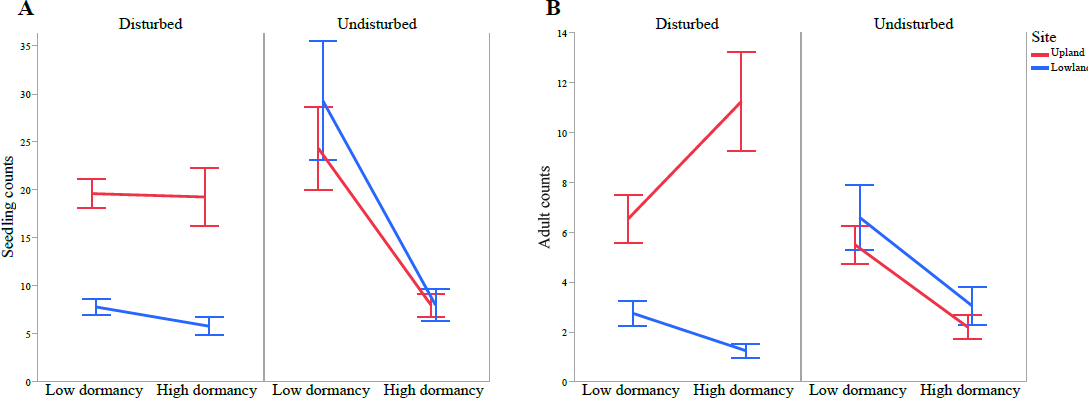
Seedling and adult recruitment for genotypes with high dormancy versus low dormancy group under disturbance treatment conditions in contrasting habitats. The color combination was chosen based on site and treatment effects. The xeric BFL site was colored dark red and the mesic NDP site was colored dark blue. The top panel separated both disturbed and undisturbed plots. The Y-axis represents the mean recruitment counts per plot and X-axis represents seed dormancy type (high dormancy vs low dormancy). Each error bar in the bar chart is constructed using ±1 standard error from the mean. A and B: seedling and adult recruitment in contrasting habitats under disturbance treatment conditions.

Furthermore, our data revealed that the low dormancy plots had significantly greater seedling recruitment compared to the high dormancy plots in both disturbed and undisturbed conditions across sites (Supplementary Table 2). However, the difference in seedling recruitment between dormancy groups was more pronounced in undisturbed plots than in plots that received disturbances. The low dormancy group exhibited 67% and 72% higher seedling recruitment in undisturbed plots compared to the high dormancy group at the BFL and NDP sites, respectively. In contrast, in disturbed plots, the low dormancy group showed a 1.5% increase (P > 0.05) and a 25% increase (P < 0.05) in seedling recruitment compared to the high dormancy group at the BFL and NDP sites, respectively (Figure 6, Supplementary Table 2).

Regarding adult recruitment, we compared disturbed and undisturbed plots for different dormancy groups at the two sites separately. Overall, an interesting pattern emerged between dormancy and disturbance. In the absence of disturbance, low dormancy plots exhibited significantly greater adult recruitment compared to high dormancy plots at both sites. However, in disturbed plots at the BFL site, high dormancy plots had significantly higher adult recruitment compared to low dormancy plots. Specifically, the high dormancy plots at BFL showed 42% more recruitment than the low dormancy plots with disturbance, while the low dormancy plots had 60% more recruitment than the high dormancy plots in the absence of disturbance. On the other hand, at the NDP site, the low dormancy plots had 54% and 53% higher adult recruitment compared to high dormancy plots in both disturbed and undisturbed plots, respectively (Figure 6, Supplementary Table 2). These results indicate that undisturbed plots favored low dormancy seeds across sites, while high dormancy seeds were favored in disturbed plots at the BFL site.

## Discussion

This study aimed to assess the influence of combining different factors, such as seed addition and surface soil disturbance, along with variation in seed mass and germination characteristics of different genotypes in diverse habitats, on the recruitment of *P. hallii*. Our investigation delves into seed quality from various angles, including the incorporation of natural genotypes and subsequent recombination of seed traits in hybrids. The findings consistently indicate intricate interactions between seed factors (abundance and potential quality) and site conditions (disturbance and habitat) as limiting factors. It is evident that the early stages of establishment significantly impact *P. hallii*. Furthermore, the results suggest that both seed size and dormancy may play a role, potentially interacting with each other. Overall, our data supports the notion of local adaptation in terms of seedling and adult recruitment and provides insights into the factors that constrain the distribution of lowland and upland ecotypes of this perennial species.

### Germination and seed bank persistence significantly varied between genotypes

Seed dormancy has posed challenges in several published studies aiming to assess recruitment, as dormant seeds can remain ungerminated throughout the study period, leading to incomplete or misleading conclusions. This phenomenon is particularly relevant in long-term ecological investigations where researchers track seedling establishment and population dynamics over extended periods. The presence of dormant seeds can artificially inflate or underestimate recruitment rates, leading to biased assessments of species’ reproductive success and population persistence. Studies have documented instances where dormant seeds germinated years or even decades after the initial seed deposition, highlighting the need for prolonged observation and accounting for seed dormancy effects Thompson et al. 1997; Merritt et al. 2007). In our experimental plots, we observed a scarcity of seedlings for both genotypes, despite sowing 100 seeds in each seed addition plot. The number of seedlings in the field was 61% lower for the lowland genotype and 17% lower for the upland genotype compared to the germination rate observed in the laboratory-based assay (Figure 3). This disparity could be attributed to potential factors such as seed predation by birds or seed-eating ants, as well as seedling pathogens that may have culled the seedlings from our experimental plots. It is also plausible that we missed some initial seedling counts that were lost due to interactions with natural enemies. However, our seed burial experiments revealed that the upland genotype exhibited greater seed bank persistence than the lowland genotype at both sites, indicating that the relatively low level of seed germination, particularly for the upland genotype, was influenced by seed dormancy and the sustained presence of supplemental seed in the seed bank (Figure 2). Despite our continuous monitoring of the field sites over multiple seasons, a considerable portion of upland genotype seeds might still remain dormant in the soil. Consequently, the scope of our conclusions based on the study period may underestimate the extent of upland recruitment.

### Genotypic divergence exhibited different patterns of recruitment across sites

Field observations across the geographic range of *P. hallii* have consistently indicated a rare co-occurrence of the lowland and upland ecotypes within natural habitats Gould et al. 2018; Lovell et al. 2018; Lowry et al. 2015; Palacio Mejía 2018; Palacio-Mejía et al. 2021). This pattern of distribution suggests the presence of seed or site limitations that may contribute to the observed spatial segregation. By conducting the experimental manipulations in this study, we discovered that the limited overlap between ecotypes is in part attributed to seed limitation. Both the representative lowland and upland genotypes exhibited higher seedling recruitment in contrasting habitats compared to control plots (Figure 3). However, the transition from the seedling stage to the adult stage revealed distinct limitations that varied depending on the genotype and site. For instance, while seedling recruitment indicated seed limitation for both genotypes, the pattern of adult recruitment suggested a more pronounced dependence on habitat, particularly for the upland genotype. The transition from seedling to adult stage for the lowland genotype was successful in both its native (mesic) site and foreign (xeric) site, whereas the upland genotype struggled to transition to the adult stage at the foreign (mesic) site. However, we observed adult recruitment of the upland genotype at the foreign (mesic) sites only when the soil surface was disturbed. This recruitment pattern was also evident at their native (xeric) site, where the upland genotype recruited more adults on disturbed plots. Conversely, the lowland genotype exhibited higher adult recruitment on undisturbed plots at both experimental sites. These recruitment patterns suggest that soil surface disturbance favored the upland genotype, while undisturbed plots favored lowland genotype recruitment across the experimental sites. These observations can be explained by the contrasting habitat structures preferred by these representative genotypes. The upland genotype typically occupies rocky, calcareous, and open canopy habitats, which aligns with their preference for less competitive environments, as evidenced by their higher recruitment on disturbed plots. In contrast, the lowland genotype thrives in sandy, high-density, competitive environments, which likely explains their preference for undisturbed plots across sites. Taken together, our findings and the observed patterns of abundance suggest that both site quality and habitat structure exert significant influences on seedling and adult recruitment in *P. hallii*.

However, the existing literature on recruitment limitations primarily focuses on seed or microsite constraints at the species or population level Münzbergová and Herben 2005). This perspective assumes, either explicitly or implicitly, that plant populations predominantly inhabit environments with homogeneous conditions, as defined by Münzbergová and Herben (2005). Nevertheless, such environmental homogeneity is more likely to be an exception rather than the norm, and this assumption holds true only in specific cases, such as certain grasslands. In the majority of plant populations, both seed availability and the factors influencing it, as well as the availability of safe sites for establishment, exhibit spatial heterogeneity and vary across habitats and microhabitats. Very few studies showed that the combined effects of seed availability and safe-site availability determine seedling recruitment dynamics at the population level Klinkhamer and De Jong 1989; Eriksson and Ehrlén 1992b; Maron and Gardner 2000; Juenger and Bergelson 2000a). Our study further highlights the variability in recruitment limitations experienced by the same species across diverse habitats. This serves as a strong reminder that characterizing individual populations as solely seed or site limited without conducting thorough investigations is inadequate. The findings underscore the importance of careful and comprehensive studies to accurately identify and understand the specific limitations influencing recruitment dynamics in different habitats.

### Seed size and dormancy trait combinations facilitate adaptation in contrasting habitats

Variations in seed size have been demonstrated to significantly influence the success of seedling establishment across diverse ecosystems (Westoby et al., 2002). The recruitment of small-seeded species is often believed to be constrained by microsite limitations due to the substantial influence of competition from dominant vegetation Harper 1977; Turnbull et al. 2000; Levine and Rees 2002). Conversely, large-seeded species are typically regarded as seed limited, as their fecundity tends to be relatively low compared to the availability of microsites Levins and Culver 1971; Shmida and Ellner 1984; Tilman 1994; Turnbull et al. 2000). Empirical evidence strongly supports the positive relationship between seed size and recruitment success Gross 1984; Burke and Grime 1996; Jakobsson and Eriksson 2000; Kidson and Westoby 2000; Turnbull et al. 2005). However, the extent to which this relationship depends on the environmental context remains unclear. In a recent study conducted with *P. hallii*, we uncovered a relatively robust genetic correlation between seed size and dormancy, with larger seeds exhibiting delayed germination Razzaque and Juenger 2022). This pattern of functional trait variation suggests that *P. hallii* ecotypes may have experienced strong selection pressures, leading to the evolution of trait combinations involving either large seed/high dormancy or small seed/low dormancy, which are associated with habitats ranging from coastal to inland sites Razzaque and Juenger 2022). Our current study revealed the significance of both traits for recruitment in xeric and mesic habitats. Contrasting the hybrids with the pure genotypes, we consistently found that parental phenotypes displayed higher recruitment in their respective native habitats, whereas the opposite trait combinations observed in hybrid lines appeared to be maladaptive. Specifically, we consistently observed that lines with larger seeds exhibited greater recruitment at the xeric site, while lines with smaller seeds displayed greater recruitment at the mesic site, aligning with the prevailing seed traits common in these regions (Figure 5). Additionally, lines with low dormancy exhibited higher seedling recruitment at both sites; however, the data on adult recruitment indicated that lines with high dormancy recruited more adults at the xeric site, whereas lines with low dormancy recruited more adults at the mesic site (Figure 6). These results suggest that combinations involving big seed/high dormancy are favored at the xeric site, while the opposite combinations are favored at the mesic site. Notably, these patterns were disrupted in the recombinant hybrids. This observation lends support to the hypothesis of local adaptation, as it demonstrates that local genotype traits perform better under their specific environmental conditions, and the selection of seed size/dormancy trait combinations is driven by the heterogeneity of habitat structure.

## Conclusion

Our study provides valuable insights into the complex dynamics of *Panicum hallii* populations, challenging the simplistic notion of strict seed or site limitation. By examining the interplay between seed and microsite availability, we reveal that the success of *P. hallii* recruitment is contingent upon both factors. Our findings highlight the significance of early establishment stages, which are intricately tied to the surrounding habitat conditions. Interestingly, our study genotypes exhibit distinct strategies for survival. The upland genotype strategically produces larger, more dormant seeds to enhance establishment in challenging, low-competition environments. Conversely, the lowland genotype opts for smaller, less dormant seeds to thrive in the highly competitive coastal prairie habitat. These observations emphasize the importance of understanding seed traits and their contribution to recruitment dynamics, ultimately enhancing our understanding of local adaptation and our ability to predict plant population outcomes. Further investigations in this field will undoubtedly yield valuable insights into the intricate mechanisms driving plant population dynamics.

## Acknowledgement

Thanks to Caroline E Farrior and Robert W Heckman for valuable discussions about experimental designs, data analysis strategies and feedback on the manuscript. Thanks to Rob Plowes from Brackenridge Field Laboratory and Jake Herring from Nueces Delta Preserve for facilitating field experiments. This research was supported by an NSF Plant Genome Research Program Grant (IOS-0922457) to TEJ. Authors acknowledge Texas Ecolab support to SR for conducting field work.

## Author contributions

SR and TEJ designed the experiment. SR conducted the experiment, collected and analyzed data and wrote the manuscript. TEJ supervised the work and edited the manuscript.

## Data Availability Statement

We have provided all raw data information in the supplementary section.

## Table and Figure legends

Supplementary Table 1: Results of generalized linear models (GLM) on seed bank longevity in contrasting habitats sampled at 18 different time points for representative lowland and upland genotypes. In the factorial model, Genotype, Site, Time and their interactions have been tested and described in the table. DF = Degrees of Freedom: L-R ChiSquare = Likelihood Ratio Chi Square. ‘*’ Indicates the significance after Bonferroni adjustment at P <0.05.

Supplementary Table 2: Results from series of planned contrasts from generalized linear model on seedling and adult recruitment. Major contrasting factors as well as specific planned contrasts were described in the table. DF = Degrees of Freedom: L-R ChiSquare = Likelihood Ratio Chi Square. ‘*’ Indicates the significance after Bonferroni adjustment at P <0.05.

Supplementary File 1: Raw seedling and adult recruitment count data for *P. hallii* in two different sites. BFL = Brackenridge Field Laboratory, Austin, Texas (xeric habitat); NDP = Nueces Delta Preserve, Odem, Texas (mesic habitat)

Supplementary File 2: Seed germination data from the seeds collected from seed burial experiment. BFL = Brackenridge Field Laboratory, Austin, Texas (xeric habitat); NDP = Nueces Delta Preserve, Odem, Texas (mesic habitat)

